# Exploring the lncRNA localization landscape within the retinal pigment epithelium under normal and stress conditions

**DOI:** 10.1101/2022.02.03.479033

**Authors:** Tadeusz J. Kaczynski, Elizabeth D. Au, Michael H. Farkas

## Abstract

Long noncoding RNAs (lncRNAs) are emerging as a class of genes whose importance has yet to be fully realized. It is becoming clear that the primary function of lncRNAs is to regulate gene expression, and they do so through a variety of mechanisms that are critically tied to their subcellular localization. Although most lncRNAs are poorly understood, mapping lncRNA subcellular localization can provide a foundation for understanding these mechanisms. Here, we present an initial step toward uncovering the localization landscape of lncRNAs in the human retinal pigmented epithelium (RPE) using high throughput RNA-Sequencing (RNA-Seq). To do this, we differentiated human induced pluripotent stem cells (iPSCs) into RPE, isolated RNA from nuclear and cytoplasmic fractions, and performed RNA-Seq on both. Furthermore, we investigated lncRNA localization changes that occur in response to oxidative stress. We discovered that, under normal conditions, most lncRNAs (76%) are seen in both the nucleus and the cytoplasm to a similar degree, but of the transcripts that are highly enriched in one compartment, more are nuclear (18.6%) than cytoplasmic (5.6%). Interestingly, under oxidative stress conditions, we observed an increase in lncRNA localization in both nuclear (23.5%) and cytoplasmic (9.7%) fractions. In addition, we found that nuclear localization was partially attributable to the presence of previously described nuclear retention motifs, while adenosine to inosine (A-to-I) RNA editing appeared to play a very minimal role. Our findings regarding lncRNA localization in the RPE provide two avenues for future research: 1) how lncRNAs function in the RPE, and 2) how one environmental factor, in isolation, may play a role in retinal disease pathogenesis.

## Introduction

lncRNAs are a poorly understood class of molecules that have garnered increased attention in recent years. Like mRNAs, lncRNAs are capped, spliced, and usually poly-adenylated, however, they only have very short open reading frames, are usually expressed at lower levels, and are more tissue-specific than their coding counterparts [1–4]. Rather than code for a protein, lncRNAs serve to modulate gene expression through transcriptional or translational control, and to do this, they must localize to either the nucleus or the cytoplasm, depending on the mechanism by which they function [5, 6].

As particularly evident in lncRNAs, RNA localization – the distribution of RNA transcripts at the subcellular level – is inextricably tied to RNA function. While our understanding of this subject is far from complete, recent publications have begun to uncover the mechanisms underlying the various aspects of RNA localization. Nuclear retention is one such aspect, wherein an RNA species is temporarily or permanently sequestered in the nucleus. Currently, nuclear retention is thought to occur via anchoring transcripts to structural entities, and/or preventing transcripts from recruiting nuclear export factors, with motifs within the primary sequence of the RNAs facilitating such processes [7]. An interrogation of the sequence of the nuclear localized *BMP2-OP1 responsive gene (Borg)* transcript identified a sequence, the BORG motif, whose presence was sufficient to impart nuclear localization, although the mechanism by which this occurred was not determined [8]. Another sequence, the 5’ splice site (5’SS) motif, identified from the backbone of an expression plasmid, promoted nuclear retention of RNAs carrying the motif through sequestration in nuclear speckles [9]. More recently, it was found that RNAs possessing a particular sequence, the SINE-derived nuclear RNA localization (SIRLOIN) motif, were bound by HNRNPK, leading to their accumulation in the nucleus [10]. Despite these discoveries, the nuclear retention of a transcript is appreciably more complex than the presence or absence of retention motifs, as demonstrated by the variable localization of such RNAs across cell types [11]. Additionally, a linear regression model based on retention motifs and other genomic and splicing features was only able to predict 15-30% of the variability in localization among lncRNAs, alluding to unknown factors contributing to RNA localization [11].

There is also a growing interest in how the phenomenon of RNA nuclear retention is used by the cell as a means of regulatory control. Indeed, recent studies have identified subsets of mRNAs which are retained in the nucleus under normal conditions and released into the cytoplasm in response to various stimuli, including: cell stress signaling [12–14], neuronal activity [15], and developmental cues [16]. The retention of these transcripts is thought to allow the cell to quickly begin synthesizing protein in response to the relevant stimulus [7]. Furthermore, research has demonstrated that nuclear retention can buffer cytoplasmic transcript levels from the noise created by bursts of transcriptional activity, thus shielding the cell from wild fluctuations in protein levels [17]. Though it is unclear whether lncRNA nuclear retention is similarly used by the cell to enact regulatory control, it seems well within the realm of possibility, given the molecular similarities between mRNAs and lncRNAs and their localization-dependent activities.

Because of its implication in regulatory processes, RNA localization is beginning to be recognized for its role in the development and progression of human disease [7]. Mutations affecting the structure and function of RNA export factors have been linked to the pathogenesis of neurodevelopmental disorders [18, 19], motoneuron diseases [20], and neurodegenerative disease [21]. Alterations in RNA primary sequence can also contribute to disease by affecting transcript localization. Repeat expansions in *HTT* and *C9ORF72* are thought to contribute to Huntington’s disease and amyotrophic lateral sclerosis (ALS), respectively, by causing the transcripts to become sequestered in the nucleus [22, 23]. Additionally, the *ApoE* transcript, a major susceptibility gene for Alzheimer’s disease (AD), has been shown to be released into the cytoplasm in response to neuronal injury in mice, implicating the dysregulation of RNA localization as a possible contributor to AD progression [24].

Age-related macular degeneration (AMD) is the third most common cause of moderate-to-severe visual impairment worldwide, currently believed to be affecting 196 million people, but the role of RNA localization in its pathogenesis is, as yet, unknown [25]. The disease results in lesions in the macula region of the eye corresponding to the death of the retinal pigmented epithelium (RPE) and the overlying photoreceptor cells [26]. Development of AMD is largely tied to environmental risk factors, with epidemiological studies finding a genetic component to account for only 37% of AMD pathogenesis [27]. Oxidative stress is a strong environmental risk factor for the progression of AMD [28]. This is attributable to the constant exposure of the retina to light, high oxygen tension, high metabolic rate, exposure to fatty acids capable of autoxidation, as well as controllable factors such as diet and smoking, which collectively contribute to the generation of reactive oxygen species in the retina and RPE [28–33]. Due to the important connection between oxidative stress in the RPE and AMD pathogenesis, several transcriptome profiling studies have examined cultured RPE exposed to oxidative stressors and native RPE from human donors with AMD [34–37]. Yet these studies have not separately examined cytoplasmic and nuclear transcriptomes, leaving unknown the subcellular distribution of RNA transcripts within the RPE cells. Thus, it is also unknown whether, how, and to what functional effect oxidative stress alters lncRNA localization in the RPE.

In this study, we examined the poly-adenylated noncoding transcriptomes of the nuclear and cytoplasmic fractions of normal human induced pluripotent stem cell derived retinal pigmented epithelium (iPSC-RPE) cells, as well as iPSC-RPE cells treated with hydrogen peroxide (H_2_O_2_) – exposure to which is commonly used to model oxidative stress in the field of AMD research [32, 33, 38, 39]. We found that, under normal conditions, the majority of lncRNAs were evenly distributed between the nucleus and cytoplasm, while the lncRNAs that localized to one fraction were overwhelmingly nuclear. However, H_2_O_2_ exposure leads more lncRNAs to commit to a subcellular compartment, with a larger proportion localizing to either the nucleus or cytoplasm. Further analysis of the data suggested that previously described nuclear retention motifs, and to a much lesser extent A-to-I RNA editing, contribute to the localization patterns under normal conditions. Yet these factors cannot explain the largescale shift in localization in response to oxidative stress, and more studies will be needed in order to understand this phenomenon.

## Results

### lncRNA Transcript Localization Shifts in Response to Oxidative Stress

We generated iPSC-RPE cells using the BXS0114 iPS cell line. One group of iPSC-RPE samples (hereafter referred to as BXS) was left untreated to act as controls, while another group of samples (hereafter referred to as BXS-H_2_O_2_) was treated with 500 μM H_2_O_2_ for 3 hours in order to induce oxidative stress. Nuclear and cytoplasmic RNA was isolated from five technical replicates of each condition, and RNA-Seq libraries were prepared. Each sample was sequenced on an Illumina NextSeq 500, and generated a minimum of 13 million reads, with at least 85% of reads uniquely aligned to the hg38 human genome build, and, on average, 92% of these reads counted. We have previously shown these data to be of high quality and the iPSC-RPE to be sufficiently RPE-like [40, 41].

We first examined overall transcript expression to assess overall sample quality and differences after exposure to H_2_O_2_. We considered a transcript to be expressed if it had an average RPKM > 0.5 across all samples in a condition. Of the 68,792 lncRNA transcripts, we found similar numbers expressed in both conditions, with 7,264 total in BXS and 7,133 BXS-H_2_O_2_. We also looked at each fraction individually to determine nuclear and cytoplasmic expression. We again see similar numbers, with more transcripts expressed in the nucleus in both conditions. Specifically, in BXS, we found 5,993 transcripts expressed in the nuclear fraction and 5,542 in the cytoplasmic fraction, while in BXS-H_2_O_2_, we found 6,294 transcripts expressed in the nucleus and 5,294 in the cytoplasm.

To examine lncRNA localization, we compared expression in the nuclear and cytoplasmic fractions. We considered a transcript to be localized if it showed at least a two-fold greater expression in one fraction, with an adjusted p-value < 0.01. A transcript with a fold-change < 1.9 between fractions, regardless of p-value, was considered to be of mixed localization - meaning that it was similarly expressed in the cytoplasmic and nuclear fractions. We found that a large portion of transcripts did not display strongly asymmetric localization, as 3,156 and 3,553 lncRNAs were of mixed localization in the control and H2O_2_-treated iPSC-RPE, respectively (Figure 1C, D). In contrast, we see far fewer transcripts localized to one fraction, with only 997 transcripts localized under control conditions. Of these, 775 were localized to the nucleus, while only 222 were cytoplasmic. Interestingly, exposure to H_2_O_2_ increases localization, as we find 1,773 transcripts localized in BXS-H_2_O_2_. We found H_2_O_2_ exposure increased localization in both fractions, with 1,253 nuclear and 520 cytoplasmic localized transcripts in the BXS-H_2_O_2_ iPSC-RPE (Figure 1C, D).

**Figure 1.**
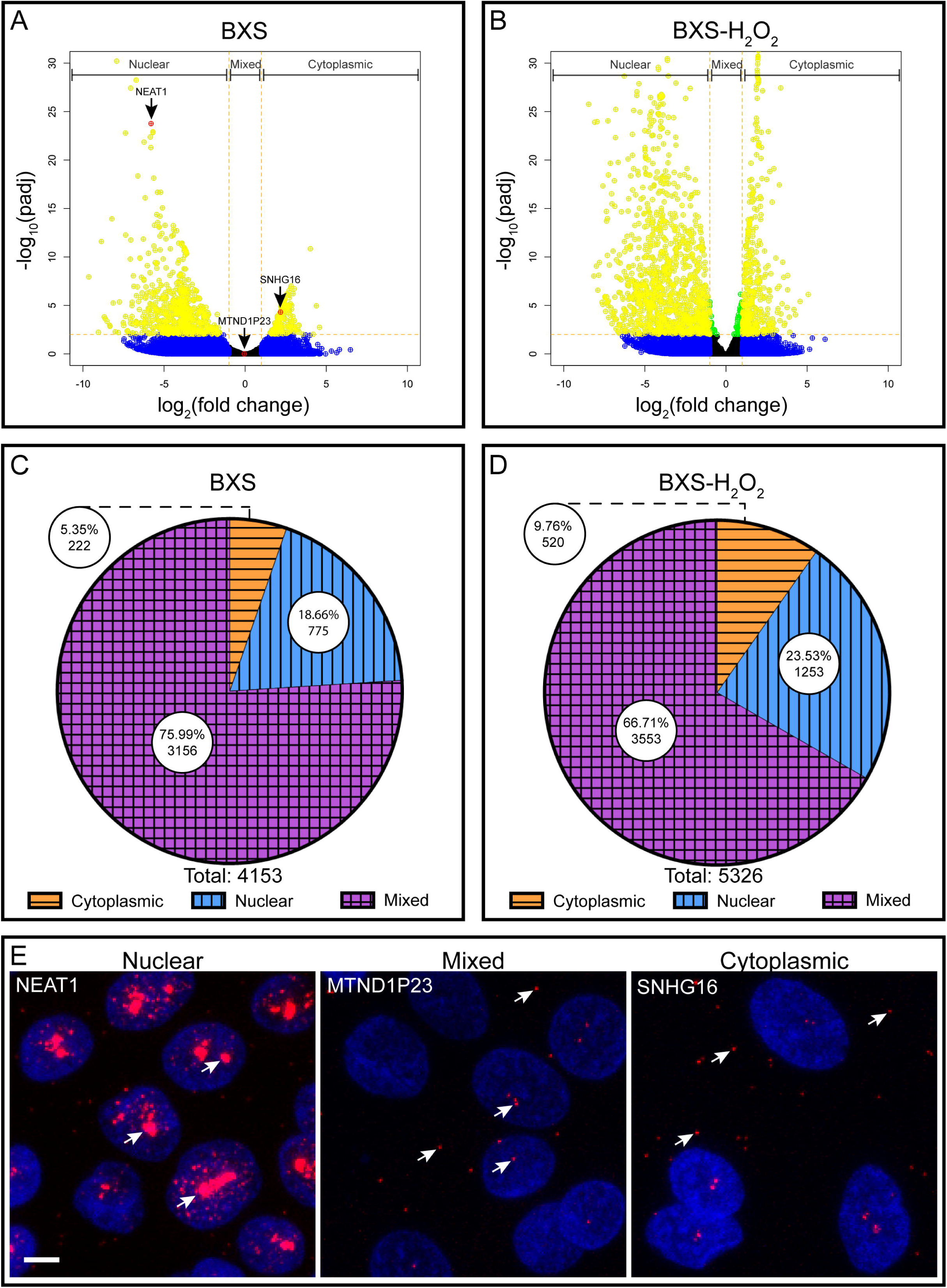
Distribution of lncRNA transcripts in the RPE. Volcano plots of lncRNAs in control BXS iPSC-RPE (A) and H_2_O_2_-treated BXS iPSC-RPE (B). Log_2_ cytoplasm:nuclear fold change and corresponding log_10_ adjusted p-value are plotted for each transcript. Transcripts with fold change >2 are colored blue, adjusted p<0.01 are green, both fold change >2 and adjusted p<0.01 are yellow. Genes confirmed via FISH are red (A). Pie graphs of the distribution of lncRNAs within the control BXS iPSC-RPE (C) and H_2_O2-treated BXS iPSC-RPE (D). The cytoplasmic categorization is indicated by orange with horizontal stripes, nuclear by blue with vertical stripes, and mixed by purple with crosshatched stripes. The total number of transcripts in each category and the percentage of the whole are indicated. (E) RNA-FISH images of iPSC-RPE confirming localization of NEAT1, MTND1P23, and SNHG16 (red) and counterstained with Hoechst solution (blue). Arrows indicate some of the localized RNAs. Scale bar is 5 μm.

In order to validate our RNA-Seq analysis, we performed RNA fluorescent in situ hybridization (RNA-FISH) using probes targeting a set of lncRNA transcripts that our analysis identified as having nuclear (NEAT1, cyto:nuc expression ratio = (<0.001)), cytoplasmic (SNHG16, cyto:nuc expression ratio = 4.9), or mixed localization (MTND1P23, cyto:nuc expression ratio = .96). We were able to confirm localization in each case (Figure 1E, S1).

### Nuclear Retention Signals Contribute to lncRNA Localization

Studies by Zhang et al. [8], Lee et al. [9], and Lubelsky and Ulitsky [10] have identified several nuclear retention motifs, however, the effects of these motifs have only been examined in small subsets of transcripts, and as such, it is unknown the extent to which such elements contribute to the overall lncRNA localization landscape. With this in mind, we set out to perform a large-scale analysis of the 5’SS, SIRLOIN, and BORG motifs to determine whether and how they influence lncRNA localization within the iPSC-RPE. Because the full SIRLOIN element is quite rare, we used the 7 nucleotide pyrimidine-rich SIRLOIN sub-element described by Lubelsky and Ulitsky [10] in our study.

Analysis of the presence one or more nuclear retention motifs in expressed lncRNA transcripts revealed a wide range of distributions for the different motifs. Whereas lncRNAs with the 5’SS motif were relatively rare (approximately 500 expressed lncRNAs), the SIRLOIN and BORG motifs were more abundant (approximately 2000 and 1700 expressed lncRNAs, respectively). In addition, we found that, similar numbers of lncRNAs containing each motif were expressed in the treated and untreated samples.

We then examined the localization of these lncRNAs in the control samples. Percentage-wise, transcripts with the 5’SS, BORG, or SIRLOIN motifs displayed similar localization patterns, with approximately 2-4% being cytoplasmic and 19-24% being nuclear (Figure 2). On the other hand, approximately 6-8% and 15-16% of lncRNAs without these motifs were localized to the cytoplasm and nucleus, respectively (Figure 2).

**Figure 2.**
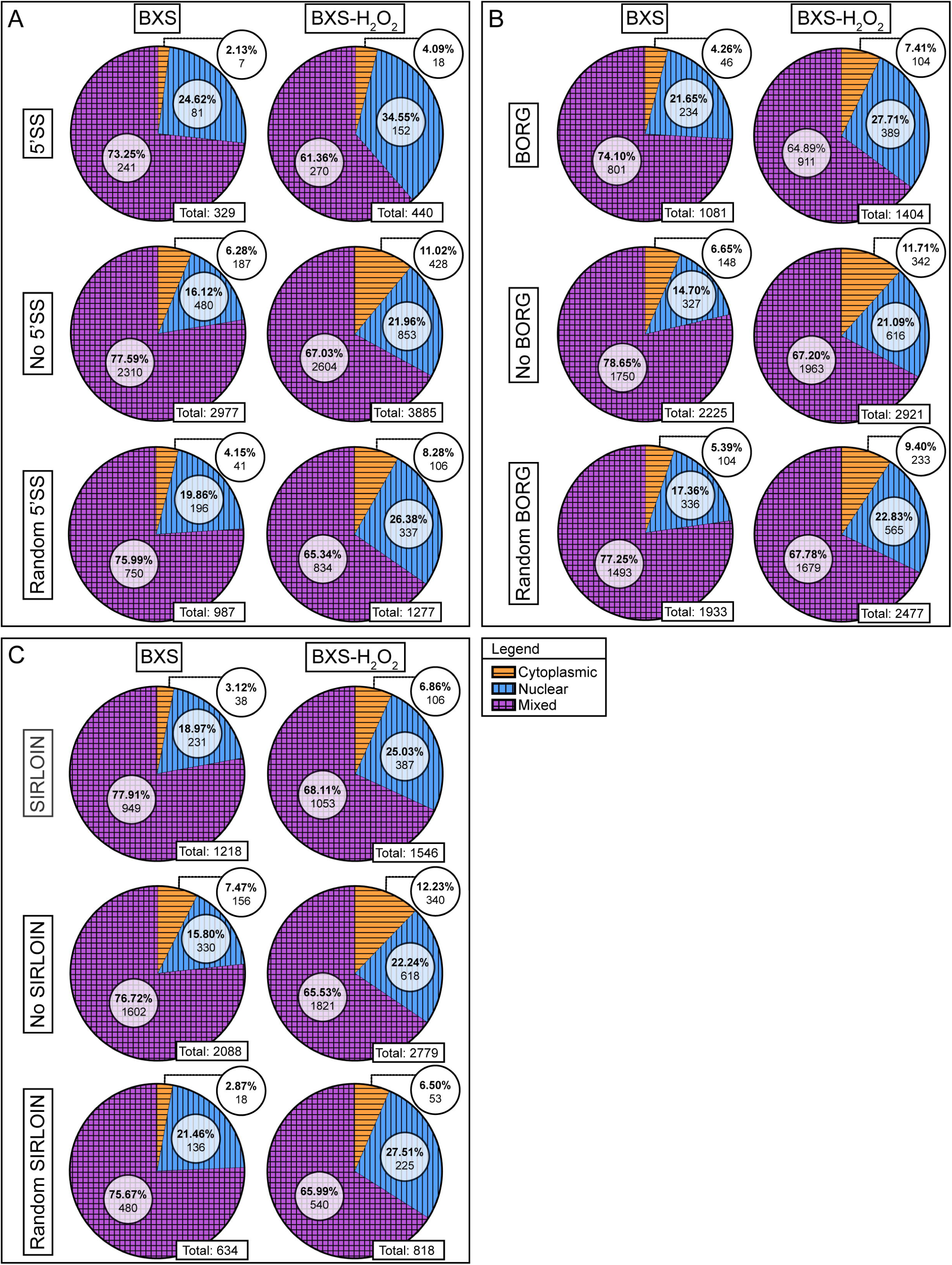
lncRNAs containing retention signal motifs are more nuclear localized. Pie graphs of the distribution of lncRNAs within the control BXS iPSC-RPE (BXS) and H_2_O_2_-treated BXS iPSC-RPE (BXS-H_2_O_2_). (A) distribution of transcripts with the 5’SS motif, without the 5’SS motif, and with a random version of the 5’SS motif. (B) distribution of transcripts with the BORG motif, without the BORG motif, and with a random version of the BORG motif. (C) distribution of transcripts with the SIRLOIN motif, without the SIRLOIN motif, and with a random version of the SIRLOIN motif. The cytoplasmic categorization is indicated by orange with horizontal stripes, nuclear by blue with vertical stripes, and mixed by purple with crosshatched stripes.

Treatment with H_2_O_2_ caused a noticeable shift in the localization patterns of ncRNAs with nuclear retention motifs. In the treated samples, approximately 11-12% and 21-22% of lncRNAs without the nuclear retention elements were localized to the cytoplasm and nucleus, respectively (Figure 2). Under oxidative stress conditions, transcripts possessing 5’SS motifs were considerably enriched in the nuclear fraction where nearly 35% of those transcripts were localized to the nucleus, contrasting the 4% which were cytoplasmic (Figure 2A). Approximately 7% and 25-27% of lncRNAs with the BORG or SIRLOIN elements were localized to the cytoplasm and nucleus, respectively, after H_2_O_2_ treatment (Figure 2B, C).

Taken together, these data indicate that under normal conditions, transcripts with 5’SS, BORG, or SIRLOIN motifs are more likely to localize to the nucleus than transcripts without such elements, yet the localization effect of oxidative stress appears to apply to all transcripts regardless of the presence or absence of these motifs. We next wanted to ensure that these localization patterns were the result of an actual phenomenon, rather than a sampling artefact from our analysis. In an attempt to decipher this, we analyzed the localization of lncRNAs possessing random sequences of the same length and structure as the 5’SS, BORG, and SIRLOIN motifs. In both the treated and untreated data sets, transcripts with random sequence variants of the SIRLOIN motifs displayed localization patterns nearly identical to transcripts with the actual SIRLOIN element (Figure 2C). On the other hand, lncRNAs with random sequence variants of the 5’SS or BORG motifs were less nuclear localized and more likely to be localized to the cytoplasm than those with the 5’SS or BORG motifs (Figure 2A, B). H_2_O_2_ exposure did not appear to affect the localization of lncRNAs with random sequence variants noticeably differently from the transcripts with the 5’SS, BORG, or SIRLOIN motifs (Figure 2).

To further our analysis, we compared transcript localization to the number of nuclear retention motifs present in each transcript. In both cell lines, and for each motif examined, we found a positive, albeit weak, correlation between the number of motifs per transcript and nuclear localization (Figure 3). In contrast, when we performed this same analysis using random motifs of 7 and 8 nucleotides, we found a mix of weak positive, negative, and neutral correlations between motif number and nuclear localization (Figure S2).

**Figure 3.**
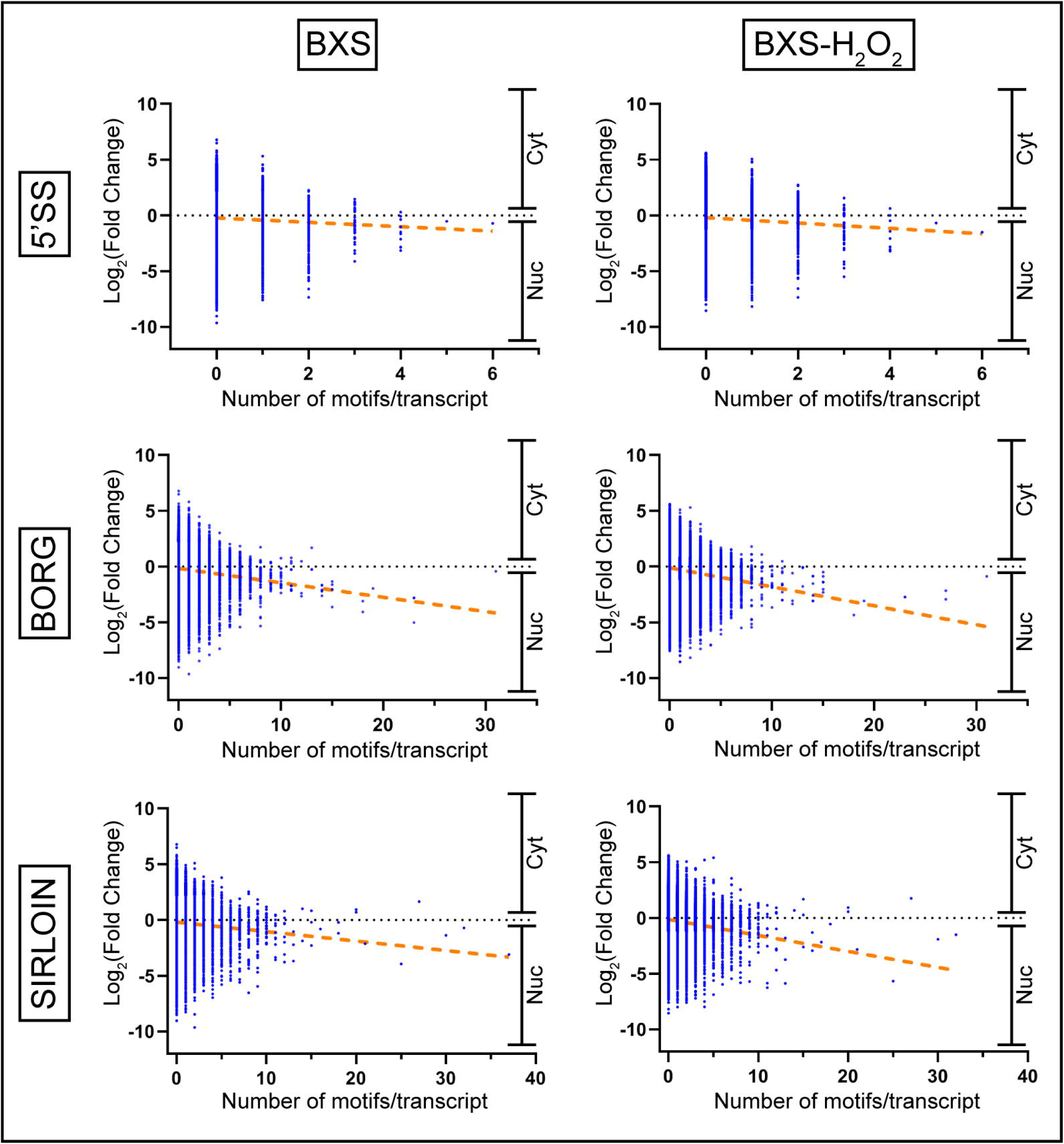
The number of retention signal motifs is positively correlated with nuclear localization. Graphs plotting the number of 5’SS, BORG, and SIRLOIN motifs per transcript versus log_2_ cytoplasm:nuclear fold change from the control BXS iPSC-RPE (BXS) and H_2_O_2_-treated BXS iPSC-RPE (BXS-H_2_O_2_) samples. Fold change corresponding to nuclear (nuc) and cytoplasmic (cyt) localization is indicated. The orange dotted lines plot the trendlines for the data.

### A-to-I RNA Editing Plays a Minimal Role in lncRNA Localization

A-to-I RNA editing has also been implicated in the nuclear retention of transcripts [14, 42, 43]. In order to better understand whether A-to-I RNA editing might be contributing to the observed localization patterns in our iPSC-RPE samples, we analyzed our RNA-Seq data for the presence of editing using SPRINT, a previously described algorithm [44]. Since unequivocal identification of editing sites within the transcriptome requires whole genome sequencing, and the genomes of the BXS0114 cell line has not yet been sequenced, our analysis was limited to the identification of potential editing sites in our samples. With the SPRINT algorithm, we identified over 200,000 potential editing sites within the transcriptomes of our samples. To probe the veracity of the analysis, ten regions containing putative editing sites identified by SPRINT were examined through Sanger sequencing – interrogating the genomic as well as transcriptomic sequences in order to discriminate edited sites from single-nucleotide polymorphisms (SNPs). Of the ten 200 bp regions examined, nine regions were found to have been edited at one or more sites (Figure 4A). The remaining region possessed a SNP at a potential editing site identified by SPRINT, but otherwise showed no evidence of editing. Of note, only 11 of the 71 editing sites detected through Sanger sequencing were also identified by SPRINT. These findings suggest that the analysis yields a low false-positive rate, yet a relatively high false-negative rate, offering a conservative estimate on the full extent of A-to-I editing.

**Figure 4.**
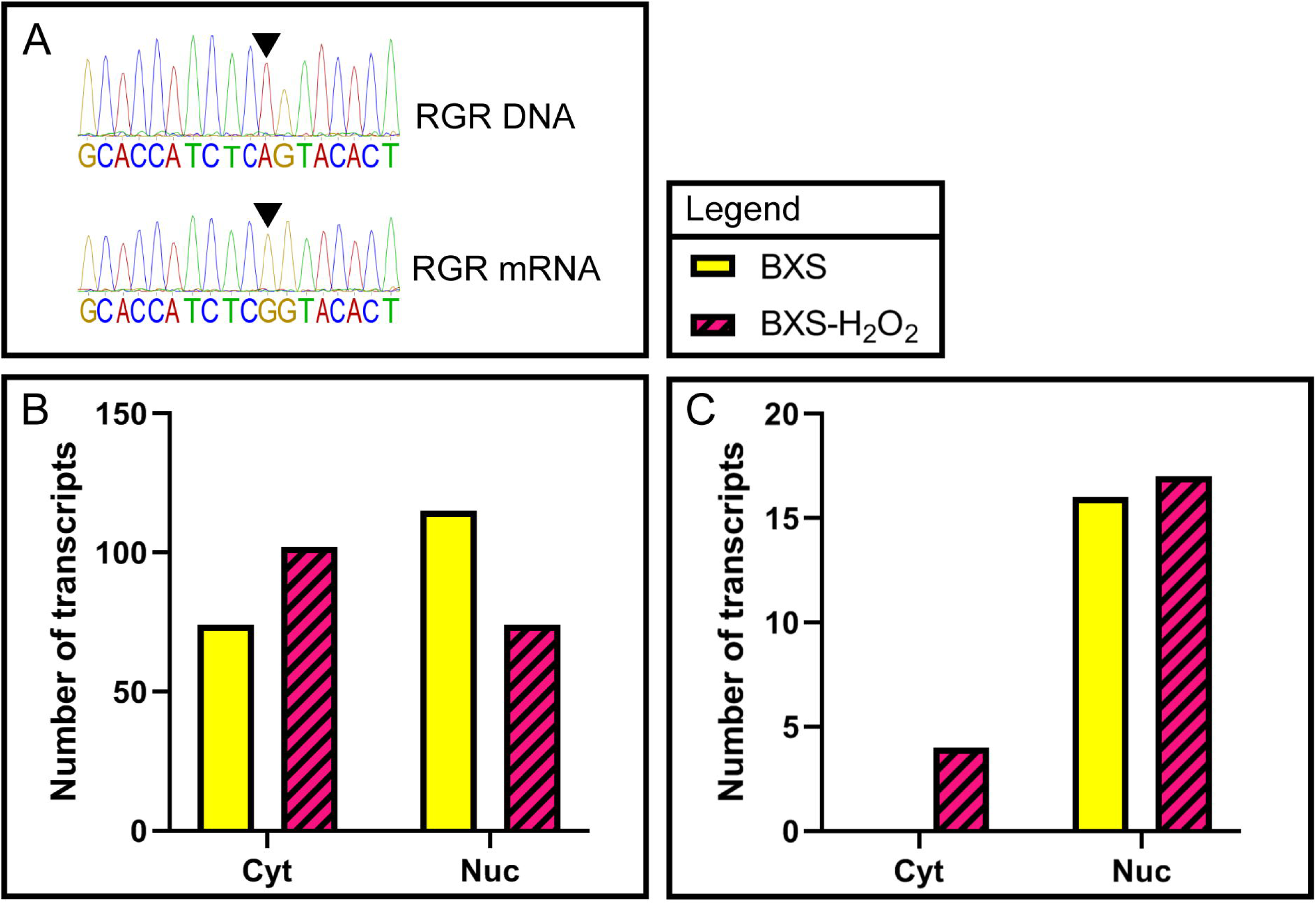
A-to-I RNA editing patterns change in response to oxidative stress. **(A)** representative sanger sequencing trace results depicting an instance of A-to-I editing within the RGR mRNA, where an inosine is read by the sequencer as a guanine. (B) Graph depicting the number of A-to-I edited lncRNAs within the cytoplasmic (cyt) and nuclear (nuc) fractions of the control BXS iPSC-RPE (BXS, solid yellow) and H_2_O_2_-treated BXS iPSC-RPE (BXS-H_2_O_2_, magenta with diagonal stripes). (C) Graph depicting the number of transcripts that were both localized to a given fraction (cytoplasmic [cyt] or nuclear [nuc]) and were more highly edited in that fraction for the control BXS iPSC-RPE (BXS) or H_2_O_2_-treated BXS iPSC-RPE (BXS-H_2_O_2_) samples.

In the untreated samples, we found 115 A-to-I edited lncRNA transcripts in the nuclear fraction and 74 edited transcripts in the cytoplasmic fraction (Figure 4B). Interestingly, in the H_2_O_2_-treated samples, we observed 74 edited lncRNA transcripts in the nuclear fraction and 102 edited transcripts in the cytoplasmic fraction – a noticeable shift in distribution compared to control (Figure 4B).

We next sought to examine the extent to which A-to-I RNA editing might contribute to nuclear retention of lncRNAs. To this end, we identified the transcripts that were both localized to a given fraction and were more highly edited in that fraction, reasoning that if A-to-I editing were driving nuclear retention, nuclear localized transcripts would be more highly edited than cytoplasmic transcripts. Indeed, in both the treated and untreated samples, we found a greater number of nuclear lncRNAs asymmetrically edited in the nucleus (BXS: 16, BXS-H_2_O_2_: 17) than cytoplasmic lncRNAs asymmetrically edited in the cytoplasm (BXS: 0, BXS-H_2_O_2_: 4) (Figure 4C).

To help contextualize the differences in editing between the H_2_O_2_-treated and untreated samples, we sought to uncover whether the expression or localization of a key A-to-I RNA editing enzyme, ADAR1-p110, changes in response to oxidative stress. A-to-I RNA editing is catalyzed by a family of enzymes called Adenosine Deaminases Acting on RNA (ADARs), and ADAR1-p110, an isoform of the ubiquitously expressed ADAR1, has been shown to translocate from the nucleus to the cytoplasm in response to UV irradiation and heat shock stresses [45, 46]. Western blotting, using an antibody that targets both the p110 and p150 isoforms of ADAR1, revealed that expression levels were not affected by H_2_O_2_ treatment (Figure S3A). Furthermore, immunofluorescent (IF) staining of ADAR1, using the same antibody, revealed no noticeable difference in ADAR1 localization between the control samples and those that were treated with peroxide (Figure S3B). To determine whether changes in ADAR1-p110 localization might be masked by co-detection with the p150 isoform, we co-transfected ARPE-19 cells with ZsGreen (a transfection marker) and a flag-tagged variant of ADAR1-p110. No noticeable translocation of flag-tagged ADAR1-p110 was observed via IF staining in response to H_2_O_2_ treatment (Figure S3C).

## Discussion

In this study, we have utilized a high-throughput approach to interrogate the localization patterns of the poly-adenylated lncRNAs within the context of iPSC-RPE. We have shown that, within untreated iPSC-RPE cells, the vast majority of lncRNA transcripts displayed a mixed localization between the nucleus and cytoplasm, and of the localized RNAs, more than 75% were nuclear (Figure 1C). These observations are largely in agreement with previously described localization patterns of lncRNAs from other human cell lines [3, 11]. It should be noted, however, that despite the tendency of lncRNAs to localize to the nucleus, the localization spectrum is wide and can vary by cell type [3]. With this in mind, and because lncRNA function is inextricably tied to subcellular localization, it is critical to understand where a transcript is localized within the relevant cell types when exploring questions of lncRNA functionality. Thus, these data will prove to be a valuable resource in understanding the functional roles played by lncRNAs in the context of the RPE.

In addition, our data corroborate the localization patterns of transcripts possessing nuclear retention motifs. Similar to the observations of Lee et al. [9], Lubelsky and Ulitsky [10], and Zhang et al. [8], we found that motif RNAs were depleted from the cytoplasmic fraction of iPSC-RPE cells (Figure 2). The localization patterns of motif RNAs were largely unchanged between the BXS and BXS-H_2_O_2_ samples, indicating that their usage was not affected by oxidative stress (Figure 2). Furthermore, our data support the finding of Zhang et al. [8] that the extent of nuclear fraction enrichment is positively correlated with the number of motifs, although we observed a much weaker correlation (Figure 2). This discrepancy likely stems from methodological differences (RT-qPCR versus RNA-Seq), and demonstrates the added nuance found using RNA-Seq analysis of subcellular fractions as a means of probing RNA localization.

Since the previously described nuclear retention motifs were present in only a subset of the nuclear localized transcripts in the iPSC-RPE cells, additional motifs or mechanisms likely exist to direct RNAs to be retained in the nucleus. Even for transcripts with known nuclear retention motifs, the molecular processes controlling the nuclear localization of RNA remain largely unclear. Though the SIRLOIN motif is thought to become bound by HNRNPK, and the 5’SS motif is believed to target RNAs to nuclear speckles, the mechanisms surrounding these processes are not yet known [9, 10]. The splicing state of a transcript is another point of consideration, as incomplete splicing may target an RNA for sequestration in the nucleus [47, 48]. Further complexity is added by the fact that nuclear retention signals may exist in RNA secondary structures such as double-stranded RNA (dsRNA) regions. Indeed, RNA is known to be involved in nucleocytoplasmic transport via interactions with proteins that specifically recognize dsRNA regions [49].

Double-stranded RNA may also be targeted for adenosine-to-inosine (A-to-I) RNA editing, which is a process that has been implicated in the nuclear retention of a number of transcripts via paraspeckle anchoring [14, 42, 43]. Our data support the notion that such RNA editing could contribute to nuclear retention, yet the relatively few A-to-I editing events revealed by our analysis make it unlikely that such a mechanism could play any more than a minor role in the distribution patterns of lncRNAs we observed in the iPSC-RPE (Figure 4B, C).

We also found that oxidative stress, in the form of H_2_O_2_ exposure, produced a dramatic shift in the lncRNA localization landscape of the iPSC-RPE cells. H_2_O_2_ treatment resulted in nearly a two-fold increase in the number of nuclear localized transcripts and a comparable fold increase in the number of cytoplasmic localized transcripts (Figure 1). In comparing the treated and untreated samples, we did not find that the increase in localized lncRNAs in the BXS-H_2_O_2_ samples could be attributed to a corresponding decrease in the number of lncRNAs with mixed localization. Nor did we find appreciable numbers of lncRNAs shifting from cytoplasmic to nuclear or vice versa. Indeed, we found the increased nuclear and cytoplasmic localization observed in the BXS-H_2_O_2_ samples to derive primarily from transcripts in the BXS samples that had fallen outside our cutoff thresholds and been left unassigned in terms of localization. Thus, the changes we observed in localization following H_2_O_2_ treatment may reflect an increased coherence between samples.

While our data indicate that oxidative insults can cause a substantial alteration in lncRNA localization within the human RPE, it is not yet clear what factors are responsible for this localization shift. Upon H_2_O_2_ treatment, lncRNAs with previously identified RNA nuclear retention motifs displayed similar shifts in localization as lncRNA transcripts that did not have those motifs (Figure 2). Additionally, while we did observe a shift in A-to-I RNA editing in lncRNAs after H_2_O_2_ exposure, very few transcripts were thus affected (Figure 4). Curiously, we saw an increase in the number of cytoplasmic A-to-I edited lncRNAs and a decrease in the number of edited nuclear lncRNAs, a phenomenon which did not appear to be caused by a translocation or change in expression of the ADAR1-p110 isoform (Figure 4B, S2). Rather than being linked to RNA localization, this increased editing may reflect a greater need for the cell to prevent the activation of the dsRNA apoptosis pathway, which is one of the known functions of ADAR1 activity [50]. Taken together, these data suggest that while the 5’SS, BORG, and SIRLOIN motifs and A-to-I RNA editing may affect RNA localization, these factors are not responsible for the major localization shift seen in the lncRNAs of the iPSC-RPE after oxidative stress.

Altered subcellular localization patterns have previously been reported in mouse cells in response to various external stimuli [12–16], but to our knowledge, this is the first examination of such changes in human iPSC-RPE cells in response to an oxidative insult and the first such study conducted at a transcriptome-wide scale. Considering that oxidative stress is a strong environmental risk factor for AMD progression [28] and that there is mounting evidence for the involvement of the dysregulation of RNA localization in the development of number of neurodegenerative diseases [21–24], these findings suggest that changes in RNA localization could play a role in the pathogenesis of AMD. Indeed, numerous lncRNAs have already been implicated in the pathology of AMD [51–54].

Much remains unknown regarding the lncRNA localization landscape within the RPE, how that localization is achieved, how it is altered by external stimuli, and how it might relate to disease pathology. Here, by mapping the localization of lncRNAs, we have set a foundation for future studies investigating the function of lncRNAs in the RPE. Further, the data presented here and in other studies, solidify the notion that transcript localization (and thus function) can vary based on cell type and environmental stressors, which serves to highlight the need to study lncRNAs in the proper context in order to understand their roles in visual dystrophies and other diseases. Future studies will enable a more thorough investigation into these remaining questions, and will not only offer insight into the roles lncRNAs may play in disease pathogenesis, but also, potentially, how they might be used for the treatment of disease.

## Materials and methods

### Culturing of cell lines and differentiation of iPSCs

All reagents were purchased from Invitrogen (Carlsbad, CA) unless noted otherwise. ARPE-19 cells (line APRE-19, ATCC, CRL-2302) were cultured in 49% Advanced DMEM (Fisher Scientific, Cat #: 12-491-015), 49% F-12 (Fisher Scientific, Cat #: MT10080CV), and 2% FBS (ATCC, Cat#: 30-2020), and were used for RNA-FISH experimentation 24-48hrs after reaching confluence. Human iPS cells (line ATCC-BXS0114, ATCC, ACS-1028) were seeded at 500,000 cells in a 10-cm dish coated with Matrigel (Fisher Scientific, Cat #: 08-774-552). Cells were maintained in TeSR-E8 media (Stem Cell Technologies, Cat #: 05990) with Rock Inhibitor (Y-27632 dihydrochloride, Santa Cruz Biotechnology, Cat #: sc-281642A) at a final concentration of 1 μM/mL. Media without Rock Inhibitor was changed daily. The procedure for differentiating human iPSCs toward RPE was performed using the BXS0114 iPSCs as previously described [40, 55] with minor adjustments. Briefly, the iPSCs were maintained until reaching 60-70% confluency, then individual colonies were lifted using Accutase (Stem Cell Technologies, Cat #: 07920). Colonies were allowed to settle in a 15 mL conical tube, old media was carefully aspirated, and fresh TeSR-E8 media was added. The colonies were then transferred to a T25 flask to initiate differentiation (day 0). Over the course of the following 4 days, the colonies were gradually transitioned to neural induction medium (NIM) (consisting of DMEM/F12, 1% N-2 supplement (Fisher Scientific, Cat #: 17-502-048), MEM non-essential amino acids (Fisher Scientific, Cat #: 11-140-050), and 2 μg/mL heparin (Sigma Aldrich, Cat #: H3149-100KU) from TeSR-E8 media in steps of 3:1, 1:1, 1:3 and 0:1 TeSR-E8:NIM. Upon reaching day 6, the colonies were transferred to a 10-cm dish coated with laminin (Fisher Scientific, Cat #: 23-017-015) in NIM, where the media was changed every 2 days. At day 16, rosettes were removed from the culture via vigorous pipetting, and the remaining cells were switched to retinal differentiation medium (consisting of DMEM/F12 (3:1), 2% B-27 supplement without retinoic acid (Fisher Scientific, Cat #: 12-587-010), and 1% antibiotic/antimycotic (Fisher Scientific, Cat #: 15-240-062)). This culture was maintained in RDM until day 80, when RPE was dissected and passaged as described. This process was performed 5 times to generate the 5 technical replicates used for this study.

### Cell fractionation

Subcellular fractionation was carried out as described by Rio et al. [56] with minor adjustments. Briefly, iPSC-RPE cells were incubated either in RDM media (untreated samples) or in RDM media with 500 μM hydrogen peroxide (treated samples) for 3 hours immediately prior to sample collection. The treatment conditions (i.e. H_2_O_2_ concentration and treatment duration) were chosen based on previous evaluations of cell death and oxidative damage analyses in ARPE-19 and human primary RPE cells in order to minimize cell death before sample collection while still sufficiently invoking oxidative stress [57–62]. Our previous analyses, indicating that iPSC-RPE and native RPE are similar from a transcriptional standpoint, support the notion that treatment outcomes would be similar in our iPSC-RPE cells [40]. The cells were then washed three times with phosphate buffered saline (PBS). Tryple Express dissociation reagent (Fisher Scientific, Cat#: 12-605-010) was applied to the cells, which were then incubated at 37°C for 5 minutes and collected via scraping. The cells were pelleted via centrifugation and resuspended in ice-cold cell disruption buffer (10 mM KCl, 1.5 mM MgCl_2_, 20 mM Tris-HCl [pH 7.5], 1 mM dithiothreitol [DTT, added just before use]). To facilitate swelling, cells were incubated on ice for 20 minutes. The cells were then transferred to an RNase-free dounce homogenizer. Homogenization was achieved using 15-20 strokes of the pestle, and the homogenate was visualized under a microscope to ensure that greater than 90% of the cell membranes were sheared while the nuclei remained intact. The homogenate was then transferred to a new 1.5 mL microcentrifuge tube. In order to strip residual cytoplasmic material from the nuclei, Triton X-100 was added to a final concentration of 0.1%, and the tubes were mixed gently by inversion. The nuclei were pelleted via centrifugation, and the supernatant (containing the cytoplasmic fraction) was transferred to a new 1.5 mL microcentrifuge tube. To wash the nuclear pellet, 1 mL of ice-cold cell disruption buffer was added, and both the nuclear and cytoplasmic fractions were centrifuged. The cytoplasmic supernatant was transferred to a new tube and the wash was removed and discarded from the nuclear pellet.

### RNA isolation

RNA was isolated from the nuclear and cytoplasmic fractions using Tri-Reagent (Molecular Research Center Inc., Cat#: TR 118) following the manufacturer’s protocol with some modifications. Briefly, after addition of the Tri-Reagent, the nuclear and cytoplasmic samples were mixed well by inversion, transferred to phase-lock heavy tubes, and incubated at room temperature for 5 minutes. 200 μL chloroform was added to each sample, which were then mixed vigorously for 15 seconds and incubated at room temperature for 15 minutes. The samples were centrifuged, and the aqueous (top) phase was transferred to a new 1.5 mL microcentrifuge tube. To remove any contaminating phenol, 400 μL chloroform was added to each sample, which were then vigorously mixed, incubated at room temperature for 2 minutes, and centrifuged. The aqueous phase, containing the RNA, was then transferred to a new 1.5 mL microcentrifuge tube. Each volume of RNA solution was then thoroughly mixed with, 1/10^th^ of a volume of 3M sodium acetate, 1 volume of isopropanol was added, and 2.5 μL RNA-grade glycogen. Precipitation of RNA was accomplished through incubation of the samples at −80°C for 1 hour. The samples were then centrifuged to pellet the RNA, and the supernatant was then discarded. To wash the RNA pellets, 75% ethanol was added to each tube, which were then briefly vortexed and centrifuged. After removal of the ethanol from the pellets, this wash step was repeated. Following the second wash, the ethanol was removed and the pellets were allowed to air dry for 5-10 minutes. The RNA was then resuspended in 22 μL DNase-free, RNase-free water and quantified via nanodrop spectrophotometer. RNA integrity and quality were assessed on a Qubit 4 Fluorometer using a Qubit RNA IQ Assay Kit. The RNA IQ values all ranged from 7.5 to 8.2, indicating high quality RNA samples.

### RNA library preparation and sequencing

RNA-sequencing libraries were prepared using the SureSelect Strand-Specific RNA Library Prep for Illumina Multiplexed Sequencing kit (Agilent, Cat#: G9691A) according to the manufacturer’s protocol. Libraries were prepared using 100 ng total RNA, and each sample was indexed for multiplexing. Prior to sequencing, library quality and quantity were determined using High Sensitivity Screen Tape on a TapeStation 4150 (Agilent). Sequencing was performed using a NextSeq 500 (Illumina) generating 2 x 150 bp reads.

### Sequencing analysis

Reads were aligned to the human genome (build hg38) with STAR v2.5.2b, and counts generated for all transcripts in gencode v29 using Rsubread v1.32.4 [63–65]. Counts were normalized using reads per kilobase per million reads (RPKM) to examine overall expression. Differential expression analysis using DESeq was performed to determine localization and effect of treatment [66]. Motif analyses were performed with gimsan. RNA editing analyses were performed using SPRINT [44].

### RNA-fluorescent in situ hybridization

iPSC-RPE cells or ARPE-19 were seeded onto 8-well chamber slides and 48-well plate coverslips, respectively, and were grown to confluence. ARPE-19 cells were chosen for their lack of pigmentation, and as a biological replicate. Cells were prepared using the ViewRNA Cell Plus Assay Kit (Fisher Scientific, Cat#: 88-19000-99) according to the manufacturer’s protocol with the minor alteration of fixation and permeabilization using 3:1 methanol:glacial acetic acid at room temperature. To stain the nuclei, the coverslips were incubated in Hoechst solution. The cells were then mounted and visualized using a Leica TCS SPE confocal microscope.

### Editing verification by sanger sequencing

Ten potential editing sites were selected from among the sites identified by the SPRINT algorithm. Primers were designed to flank the selected sites in order to produce amplicons of approximately 200 bp. PCR amplification of these sites was performed using genomic DNA isolated from BXS0114 iPSC-RPE cells and using cDNA synthesized from RNA isolated from BXS0114 iPSC-RPE cells. Following PCR amplification and gel electrophoresis, the amplicons were excised from the gel and purified. The purified amplicons were cloned into the pCR-4Blunt-TOPO vector (Fisher Scientific, Cat #: 45-003-1), the plasmids were isolated, and the inserts were Sanger sequenced using M13 Forward primers. Alignment of the genomic and transcriptomic sequences parsed bona fide editing sites from SNPs.

### Immunoblotting analysis

ARPE-19 cells were incubated either in media (untreated samples) or in media with 500 μM hydrogen peroxide (treated samples) for 3 hours immediately prior to sample collection. Cell lysates were collected using 1x RIPA buffer (Abcam, Cat #: ab156034), and protein concentrations were determined using a BCA protein assay (Fisher Scientific, Cat #: PI23227). The samples were electrophoresed on a 4-12% Bis-Tris protein gel (Fisher Scientific, Cat #: NP0335BOX), and proteins were then transferred to a nitrocellulose membrane (Fisher Scientific, Cat #: 45-004-012). The membranes were blocked with 1x Tris buffered saline with 1% casein (Bio-Rad, Cat #: 1610782) for 1 hour at room temperature, and incubated with either anti-ADAR1 (Abcam, Cat #: ab88574) or anti-tubulin (Novus Biologicals, Cat #: NB100-690) primary antibodies overnight at 4°C. Then, the membranes were washed with 1x Tris-buffered saline with 0.1% Tween 20 (TBST), incubated with an alkaline phosphatase conjugated secondary antibody (Krackeler Scientific, 45-A4312-.25mL) for 1 hour at room temperature, and developed using ECF substrate (Fisher Scientific, Cat #: 45-000-947).

### ADAR1 transfection and immunofluorescence

ZsGreen and flag-tagged ADAR1-p110 mRNA were made using the Invitrogen MEGAscript T7 Transcription Kit (Fisher Scientific, Cat #: AM1334) according to the manufacturer’s protocol. ARPE-19 cells were transfected with these mRNAs using the Invitrogen Lipofectamine MessengerMAX Transfection Reagent (Fisher Scientific, Cat #: LMRNA001) according to the manufacturer’s protocol. ARPE-19 cells were incubated either in media (untreated samples) or in media with 500 μM hydrogen peroxide (treated samples) for 3 hours immediately prior to sample collection. For transfected samples, hydrogen peroxide treatment was carried out 24 hours post-transfection. Immunofluorescent staining was carried out as follows: cells were washed with 1x phosphate buffered saline (PBS), fixed with ice-cold methanol for 1 minute, permeabilized with 0.2% Triton X-100 (in 1x PBS) at room temperature for 10 minutes, blocked with blocking buffer (1% bovine serum albumin, 0.03% Triton X-100, in 1x PBS) for 1 hour at room temperature, incubated in primary antibody (either anti-ADAR1 [Abcam, Cat #: ab88574] or anti-flag [Krackeler Scientific, Cat #: 45-F3165-1MG]) for 1 hour at room temperature, incubated in secondary antibody (either Alexa Fluor 488 goat anti-mouse [Fisher Scientific, Cat #: A11029] or Alexa Fluor 555 donkey anti-mouse [Fisher Scientific, Cat #: A31570]) for 1 hour at room temperature, counterstained with Hoechst (Fisher Scientific, Cat#: H3570), and then mounted on slides for visualization using a Leica TCS SPE confocal microscope.

## Author Contributions

TJK and MHF designed the study. TJK and MHF performed experiments. EDA and MHF analyzed and interpreted the data. MHF, EDA, and TJK wrote and edited the manuscript.

## Data Availability Statement

The datasets generated for this study can be found in the GEO REPOSITORY, GEO accession GSE158909.

## Acknowledgments

Facilities and resources provided by the VA Western New York Healthcare System to MHF. MHF is also a Research Biologist at the VA Western New York Healthcare System, Buffalo, NY. Computational support was provided by the Center for Computational Research at the University at Buffalo. Next Generation Sequencing services were provided by the Genomics Core at the Children’s Hospital Los Angeles. The contents of this manuscript do not reflect those of the Department of Veterans Affairs or the U.S. Government.

---

Supplementary Figure 1. Verification of lncRNA localization in ARPE-19 cells. RNA-FISH images of ARPE-19 cells confirming localization of NEAT1, MTND1P23, and SNHG16 (red) and counterstained with Hoechst solution (blue). Arrows indicate some of the localized RNAs. Scale bar is 5 μm.

Supplementary Figure 2. The number of random motifs is not correlated well with localization. Graphs plotting the number of 7-nucleotide (A) and 8-nucleotide (B) motifs per transcript versus log_2_ cytoplasm:nuclear fold change from the control BXS iPSC-RPE (BXS) and H_2_O_2_-treated BXS iPSC-RPE (BXS-H_2_O_2_) samples. Fold change corresponding to nuclear (nuc) and cytoplasmic (cyt) localization is indicated. The orange dotted lines plot the trendlines for the data. Five different random motifs of each length were analyzed.

Supplementary Figure 3. Analysis of ADAR1 expression and localization. (A) Detection of ADAR1 by western blotting of lysates from ARPE-19 cells that were treated with H_2_O_2_ or left untreated. (B) Immunofluorescent staining of ADAR1 in ARPE-19 cells that were treated with H2O_2_ or left untreated. (C) Immunofluorescent detection of ADAR1-p110 in ARPE-19 cells that were treated with H_2_O_2_ or left untreated and that were transfected with either only ZsGreen mRNA (Neg. Control) or ZsGreen mRNA plus ADAR1-p110-Flag mRNA (ADAR1-p110). Successful transfection was visualized by fluorescence of the ZsGreen protein, and ADAR1-p110 expression was visualized via an anti-flag antibody.

## References

1. Mercer, T.R., M.E. Dinger, and J.S. Mattick, Long non-coding RNAs: insights into functions. Nature Reviews Genetics, 2009. 10(3): p. 155–159.

2. Cabili, M.N., et al., Integrative annotation of human large intergenic noncoding RNAs reveals global properties and specific subclasses. Genes & Development, 2011. 25(18): p. 1915–1927.

3. Derrien, T., et al., The GENCODE v7 catalog of human long noncoding RNAs: Analysis of their gene structure, evolution, and expression. Genome Research, 2012. 22(9): p. 1775–1789.

4. Kashi, K., et al., Discovery and functional analysis of lncRNAs: Methodologies to investigate an uncharacterized transcriptome. Biochimica Et Biophysica Acta-Gene Regulatory Mechanisms, 2016. 1859(1): p. 3–15.

5. Bridges, M.C., A.C. Daulagala, and A. Kourtidis, LNCcation: lncRNA localization and function. Journal of Cell Biology, 2021. 220(2).

6. Marchese, F.P., I. Raimondi, and M. Huarte, The multidimensional mechanisms of long noncoding RNA function. Genome Biology, 2017. 18.

7. Wegener, M. and M. Muller-McNicoll, Nuclear retention of mRNAs - quality control, gene regulation and human disease. Semin Cell Dev Biol, 2018. 79: p. 131–142.

8. Zhang, B., et al., A novel RNA motif mediates the strict nuclear localization of a long noncoding RNA. Mol Cell Biol, 2014. 34(12): p. 2318–29.

9. Lee, E.S., et al., The consensus 5′ splice site motif inhibits mRNA nuclear export. PLoS One, 2015. 10(3): p. e0122743.

10. Lubelsky, Y. and I. Ulitsky, Sequences enriched in Alu repeats drive nuclear localization of long RNAs in human cells. Nature, 2018. 555(7694): p. 107–111.

11. Zuckerman, B. and I. Ulitsky, Predictive models of subcellular localization of long RNAs. Rna, 2019. 25(5): p. 557–572.

12. Boutz, P.L., A. Bhutkar, and P.A. Sharp, Detained introns are a novel, widespread class of post-transcriptionally spliced introns. Genes Dev, 2015. 29(1): p. 63–80.

13. Ninomiya, K., N. Kataoka, and M. Hagiwara, Stress-responsive maturation of Clk1/4 pre-mRNAs promotes phosphorylation of SR splicing factor. J Cell Biol, 2011. 195(1): p. 27–40.

14. Prasanth, K.V., et al., Regulating gene expression through RNA nuclear retention. Cell, 2005. 123(2): p. 249–63.

15. Mauger, O., F. Lemoine, and P. Scheiffele, Targeted Intron Retention and Excision for Rapid Gene Regulation in Response to Neuronal Activity. Neuron, 2016. 92(6): p. 1266–1278.

16. Yap, K., et al., Coordinated regulation of neuronal mRNA steady-state levels through developmentally controlled intron retention. Genes Dev, 2012. 26(11): p. 1209–23.

17. Bahar Halpern, K., et al., Nuclear Retention of mRNA in Mammalian Tissues. Cell Rep, 2015. 13(12): p. 2653–62.

18. Amos, J.S., et al., Autosomal recessive mutations in THOC6 cause intellectual disability: syndrome delineation requiring forward and reverse phenotyping. Clin Genet, 2017. 91(1): p. 92–99.

19. Kumar, R., et al., THOC2 Mutations Implicate mRNA-Export Pathway in X-Linked Intellectual Disability. Am J Hum Genet, 2015. 97(2): p. 302–10.

20. Nousiainen, H.O., et al., Mutations in mRNA export mediator GLE1 result in a fetal motoneuron disease. Nat Genet, 2008. 40(2): p. 155–7.

21. Ylikallio, E., et al., MCM3AP in recessive Charcot-Marie-Tooth neuropathy and mild intellectual disability. Brain, 2017. 140(8): p. 2093–2103.

22. de Mezer, M., et al., Mutant CAG repeats of Huntingtin transcript fold into hairpins, form nuclear foci and are targets for RNA interference. Nucleic Acids Research, 2011. 39(9): p. 3852–3863.

23. Freibaum, B.D., et al., GGGGCC repeat expansion in C9orf72 compromises nucleocytoplasmic transport. Nature, 2015. 525(7567): p. 129–33.

24. Xu, Q., et al., Intron-3 retention/splicing controls neuronal expression of apolipoprotein E in the CNS. J Neurosci, 2008. 28(6): p. 1452–9.

25. Jonas, J.B., C.M.G. Cheung, and S. Panda-Jonas, Updates on the Epidemiology of Age-Related Macular Degeneration. Asia Pac J Ophthalmol (Phila), 2017. 6(6): p. 493–497.

26. Hernandez-Zimbron, L.F., et al., Age-Related Macular Degeneration: New Paradigms for Treatment and Management of AMD. Oxid Med Cell Longev, 2018. 2018: p. 8374647.

27. DeAngelis, M.M., et al., Genetics of Age-Related Macular Degeneration: Current Concepts, Future Directions. Seminars in Ophthalmology, 2011. 26(3): p. 77–93.

28. Cano, M., et al., Cigarette smoking, oxidative stress, the anti-oxidant response through Nrf2 signaling, and Age-related Macular Degeneration. Vision Res, 2010. 50(7): p. 652–64.

29. Beatty, S., et al., The role of oxidative stress in the pathogenesis of age-related macular degeneration. Surv Ophthalmol, 2000. 45(2): p. 115–34.

30. Datta, S., et al., The impact of oxidative stress and inflammation on RPE degeneration in non-neovascular AMD. Prog Retin Eye Res, 2017. 60: p. 201–218.

31. Jarrett, S.G. and M.E. Boulton, Consequences of oxidative stress in age-related macular degeneration. Mol Aspects Med, 2012. 33(4): p. 399–417.

32. Jin, G.F., J.S. Hurst, and B.F. Godley, Hydrogen peroxide stimulates apoptosis in cultured human retinal pigment epithelial cells. Current Eye Research, 2001. 22(3): p. 165–173.

33. Lu, L., et al., Effects of different types of oxidative stress in RPE cells. J Cell Physiol, 2006. 206(1): p. 119–25.

34. Donato, L., et al., miRNAexpression profile of retinal pigment epithelial cells under oxidative stress conditions. FEBS Open Bio, 2018. 8(2): p. 219–233.

35. Kim, E.J., et al., Complete Transcriptome Profiling of Normal and Age-Related Macular Degeneration Eye Tissues Reveals Dysregulation of Anti-Sense Transcription. Sci Rep, 2018. 8(1): p. 3040.

36. Newman, A.M., et al., Systems-level analysis of age-related macular degeneration reveals global biomarkers and phenotype-specific functional networks. Genome Med, 2012. 4(2): p. 16.

37. Voigt, A.P., et al., Single-cell transcriptomics of the human retinal pigment epithelium and choroid in health and macular degeneration. Proc Natl Acad Sci U S A, 2019. 116(48): p. 24100–24107.

38. Bailey, T.A., et al., Oxidative stress affects the junctional integrity of retinal pigment epithelial cells. Investigative Ophthalmology & Visual Science, 2004. 45(2): p. 675–684.

39. Szaflik, J.P., et al., DNA damage and repair in age-related macular degeneration. Mutation Research-Fundamental and Molecular Mechanisms of Mutagenesis, 2009. 669(1-2): p. 169–176.

40. Au, E.D., et al., Characterization of lincRNA expression in the human retinal pigment epithelium and differentiated induced pluripotent stem cells. PLoS One, 2017. 12(8): p. e0183939.

41. Kaczynski, T.J., E.D. Au, and M.H. Farkas, Oxidative stress alters transcript localization of disease-causing genes in the retinal pigment epithelium. bioRxiv, 2021: p. 2021.01.07.425741.

42. Chen, L.L. and G.G. Carmichael, Altered nuclear retention of mRNAs containing inverted repeats in human embryonic stem cells: functional role of a nuclear noncoding RNA. Mol Cell, 2009. 35(4): p. 467–78.

43. Zhang, Z. and G.G. Carmichael, The fate of dsRNA in the nucleus: a p54(nrb)-containing complex mediates the nuclear retention of promiscuously A-to-I edited RNAs. Cell, 2001. 106(4): p. 465–75.

44. Zhang, F., et al., SPRINT: an SNP-free toolkit for identifying RNA editing sites. Bioinformatics, 2017. 33(22): p. 3538–3548.

45. Nishikura, K., A-to-I editing of coding and non-coding RNAs by ADARs. Nat Rev Mol Cell Biol, 2016. 17(2): p. 83–96.

46. Sakurai, M., et al., ADAR1 controls apoptosis of stressed cells by inhibiting Staufen1-mediated mRNA decay. Nature Structural & Molecular Biology, 2017. 24(6): p. 534-+.

47. Dias, A.P., et al., A role for TREX components in the release of spliced mRNA from nuclear speckle domains. Nature Communications, 2010. 1.

48. Girard, C., et al., Post-transcriptional spliceosomes are retained in nuclear speckles until splicing completion. Nat Commun, 2012. 3: p. 994.

49. Banerjee, S. and P. Barraud, Functions of double-stranded RNA-binding domains in nucleocytoplasmic transport. Rna Biology, 2014. 11(10): p. 1226–1232.

50. Liddicoat, B.J., et al., RNA EDITING RNA editing by ADAR1 prevents MDA5 sensing of endogenous dsRNA as nonself. Science, 2015. 349(6252): p. 1115–1120.

51. Chen, Q., et al., Pathogenic Role of microRNA-21 in Diabetic Retinopathy Through Downregulation of PPARalpha. Diabetes, 2017. 66(6): p. 1671–1682.

52. Kutty, R.K., et al., Proinflammatory cytokine interferon-gamma increases the expression of BANCR, a long non-coding RNA, in retinal pigment epithelial cells. Cytokine, 2018. 104: p. 147–150.

53. Xu, X.D., et al., Long non-coding RNAs: new players in ocular neovascularization. Mol Biol Rep, 2014. 41(7): p. 4493–505.

54. Zhu, W., et al., Identification of lncRNAs involved in biological regulation in early age-related macular degeneration. Int J Nanomedicine, 2017. 12: p. 7589–7602.

55. Gamm, D.M. and J.S. Meyer, Directed differentiation of human induced pluripotent stem cells: a retina perspective. Regenerative Medicine, 2010. 5(3): p. 315–317.

56. Rio, D.C., et al., Preparation of cytoplasmic and nuclear RNA from tissue culture cells. Cold Spring Harb Protoc, 2010. 2010(6): p. pdb prot5441.

57. Ferrington, D.A., et al., Altered bioenergetics and enhanced resistance to oxidative stress in human retinal pigment epithelial cells from donors with age-related macular degeneration. Redox Biology, 2017. 13: p. 255–265.

58. Golestaneh, N., et al., Dysfunctional autophagy in RPE, a contributing factor in age-related macular degeneration. Cell Death & Disease, 2017. 8.

59. Koller, A., et al., Cysteinyl leukotriene receptor 1 modulates autophagic activity in retinal pigment epithelial cells. Scientific Reports, 2020. 10(1).

60. Mitter, S.K., et al., Dysregulated autophagy in the RPE is associated with increased susceptibility to oxidative stress and AMD. Autophagy, 2014. 10(11): p. 1989–2005.

61. Terluk, M.R., et al., N-Acetyl-L-cysteine Protects Human Retinal Pigment Epithelial Cells from Oxidative Damage: Implications for Age-Related Macular Degeneration. Oxidative Medicine and Cellular Longevity, 2019. 2019.

62. Tohari, A.M., et al., Vitamin D Attenuates Oxidative Damage and Inflammation in Retinal Pigment Epithelial Cells. Antioxidants, 2019. 8(9).

63. Dobin, A., et al., STAR: ultrafast universal RNA-seq aligner. Bioinformatics, 2013. 29(1): p. 15–21.

64. Harrow, J., et al., GENCODE: the reference human genome annotation for The ENCODE Project. Genome Res, 2012. 22(9): p. 1760–74.

65. Liao, Y., G.K. Smyth, and W. Shi, The R package Rsubread is easier, faster, cheaper and better for alignment and quantification of RNA sequencing reads. Nucleic Acids Research, 2019. 47(8).

66. Anders, S.H., W., Differential expression of RNA-Seq data at the gene level–the DESeq package. Heidelberg, Germany: European Molecular Biology Laboratory (EMBL), 2012.

