## Supplementary figures and images for "Exploring the lncRNA localization landscape within the retinal pigment epithelium under normal and stress conditions"

### Supplemental Figure 1

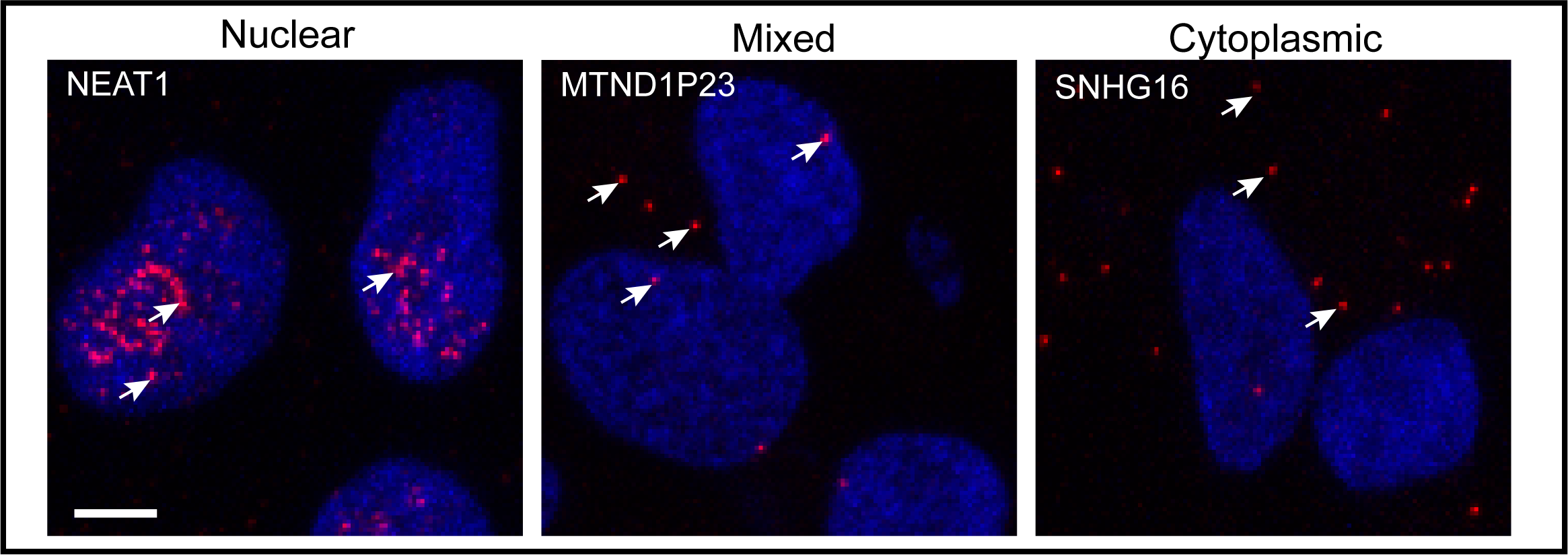

### Supplemental Figure 2

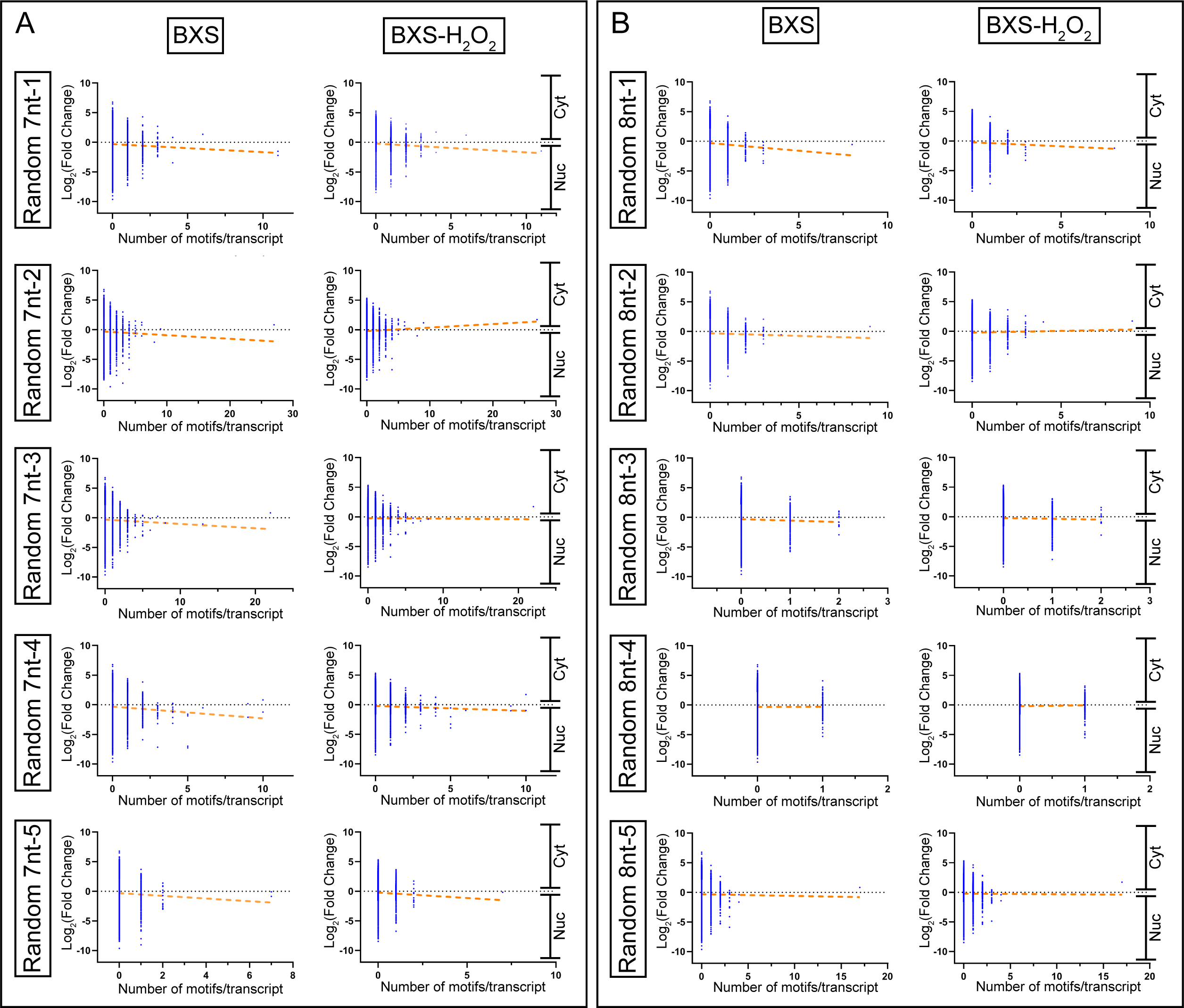

### Supplemental Figure 3

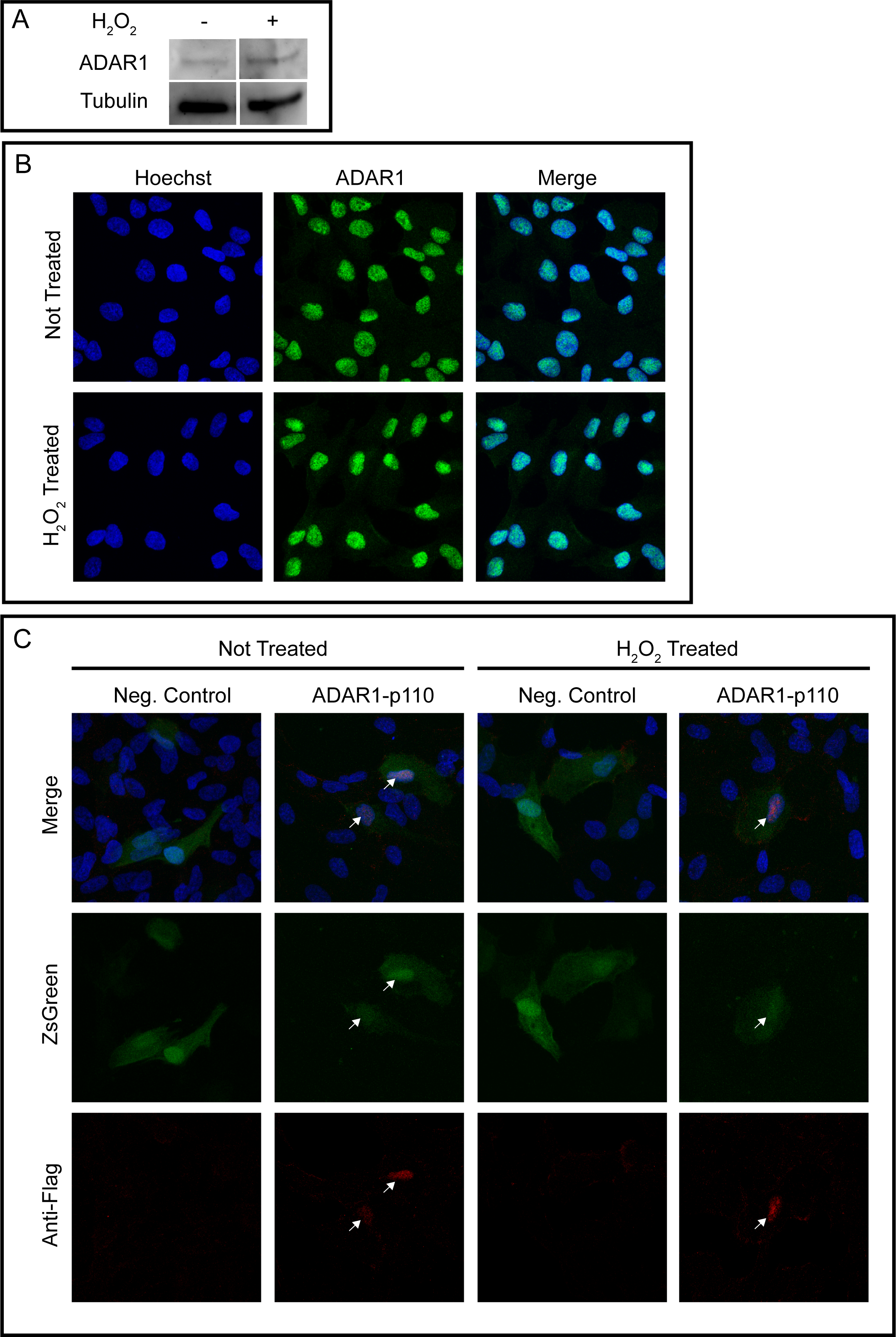
